# *Serratia marcescens* secretes proteases and chitinases with larvicidal activity against *Anopheles dirus*

**DOI:** 10.1101/2020.05.31.123539

**Authors:** Natapong Jupatanakul, Jutharat Pengon, Shiela Marie Gines Selisana, Waeowalee Choksawangkarn, Nongluck Jaito, Atiporn Saeung, Ratchanu Bunyong, Navaporn Posayapisit, Khrongkhwan Thammatinna, Nuttiya Kalpongnukul, Kittipat Aupalee, Trairak Pisitkun, Sumalee Kamchonwongpaisan

## Abstract

Vector control, the most efficient tool to reduce mosquito-borne disease transmission, has been compromised by the rise of insecticide resistance. Recent studies suggest the potential of mosquito-associated microbiota as a source for new biocontrol agents or new insecticidal chemotypes. In this study, we identified a strain of *Serratia marcescens* that has larvicidal activity against *Anopheles dirus*, an important malaria vector in Southeast Asia. This bacterium secretes heat-labile larvicidal macromolecules when cultured under static condition at 25°C but not 37°C. Two major protein bands of approximately 55 kDa and 110 kDa were present in spent medium cultured at 25°C but not at 37°C. The Liquid Chromatography-Mass Spectrometry (LC-MS) analyses of these two protein bands identified several proteases and chitinases that were previously reported for insecticidal properties against agricultural insect pests. The treatment with protease and chitinase inhibitors led to a reduction in larvicidal activity, confirming that these two groups of enzymes are responsible for the macromolecule’s toxicity. Taken together, our results suggest a potential use of these enzymes in the development of larvicidal agents against *Anopheles* mosquitoes.

## 1. Introduction

Aside from nuisances, mosquitoes are vectors of a wide range of human pathogens including *Plasmodium* parasites that cause malaria, filarial nematodes and viruses such as dengue, Zika, yellow fever, and chikungunya. These pathogens cause diseases that are major public health and economical burdens causing approximately 500,000 deaths each year (Shaw and Catteruccia, 2019), with malaria alone accounting for 90% of the deaths. *Anopheles dirus* has been incriminated as the primary vector of the malarial protozoa *P. falciparum* and *P. vivax* in Southeast Asia (Manguin et al., 2008). Although malaria cases and disease burden have significantly decreased over the past decades owing to an extensive effort in mosquito control and antimalarial drug treatment, malaria elimination faces obstacles such as the increasing insecticide resistance of mosquitoes as well as the emergence of antimalarial drug resistance of *Plasmodium* parasites (Whitty and Ansah, 2019). While there are several new classes of antimalarial drugs in the clinical pipelines (“MMV-supported projects | Medicines for Malaria Venture,” 2020; Wells et al., 2015), insecticides with new mode of action are much less developed (Shaw and Catteruccia, 2019; Whitty and Ansah, 2019).

Mosquitoes live in constant interactions with microorganisms in their environment and throughout their life cycle (Gendrin and Christophides, 2013; Wang et al., 2011), and in particular, the aquatic stages of the mosquito larvae because they are submerged in water (Dickson et al., 2017). Although microbiota are needed for the healthy development of the mosquito larva (Chouaia et al., 2012; Coon et al., 2014; Dickson et al., 2017; Valzania et al., 2018), certain microbes can be pathogenic to them (Baumann et al., 1991; Laurence et al., 2011). These entomopathogenic microorganisms, such as *Bacillus thuringiensis* and *Bacillus sphaericus* or their by-products, have long been used for mosquito population control (Baumann et al., 1991; Laurence et al., 2011). In addition to these *Bacillus* species, recent studies have identified entomopathogenic bacteria from the mosquito gut microbiota such as *Chromobacterium* spp. and *Serratia marcescens* (Bahia et al., 2014; Ramirez et al., 2014; Short et al., 2018). *Chromobacterium* spp. isolated from the mosquito gut from Panama *(Csp_P)* has insecticidal properties and has potential to be developed as a mosquito larvicidal agent (Caragata et al., 2020). This suggests that the natural microbial communities associated with mosquitoes are invaluable resources for the discovery of biocontrol agents with new mode of actions and the development of novel insecticides.

Our laboratory has recently set up a new insectary facility to expand our drug development pipeline to test antimalarial compounds against sexual stage of *Plasmodium* parasites. During the optimization of *Anopheles dirus* rearing condition, we found that if the larval food particle size was not fine enough, the food sedimented to the bottom of the pan. Due to the surface feeding nature of *An. dirus*, the sedimented food led to a bacterial growth and biofilm at the bottom of the pan. Mortality of the mosquito larva was observed a few days after the biofilm formed at the bottom of the pan. From this observation, we hypothesized that certain microbes in the larval rearing pan secrete molecule(s) that have larvicidal activity against *An. dirus* after the bacteria grow to a certain density when culture under static condition. In this study, we isolated and identified culturable bacteria with distinct morphology from the *An. dirus* larval rearing pan where larval death was being observed. Of seven isolates, we found a strain of *Serratia marcescens* that secreted heat-labile larvicidal macromolecules into spent medium when it was cultured under static condition at 25°C. We then used mass spectrometry to identify chitinases and proteases as potential larvicidal molecules. Chitinase and protease inhibitors were then used to confirm that these molecules contributed to larvicidal activity. This study demonstrates that the chitinases and proteases uncovered in this study could potentially be used as larvicides against *Anopheles* mosquito.

## 2. Materials and Methods

### 2.1 Ethics statement

Animal work in this study was carried out in accordance to the recommendations provided in the Guide for the Care and Use of Laboratory Animals of the National Institutes of Health. Mice were used only as a blood source for mosquito rearing, according to the approved protocol (BT-Animal 26/2560). Larvicidal activity assays were performed following the approved protocol (BT-Animal 20/2560). Both animal protocols were approved by the BIOTEC Committee for use and care of laboratory animals.

### 2.2 Mosquito rearing condition

Specimens of laboratory strain of *An. dirus* were originally collected in Mae Sod District, Tak Province, Thailand. A colony of *An. dirus* was maintained at the Department of Parasitology, Chiang Mai University and BIOTEC’s insectary at 27 °C with 75% humidity and a 12-h day/night, 30-min dusk/ dawn lighting cycle. The larvae were fed a diet of powdered fish food (Tetrabit, Germany). Adults were fed on a 10% sucrose solution supplemented with 5% multivitamin syrup (Sevenseas, UK) *ad libitum* (Choochote and Saeung, 2013). Female ICR mice were anesthetized using 2% Tribromoethanol and used as a blood source to maintain mosquito colony (approved protocol BT-Animal 26/2560)

### 2.3 Bacterial isolation and identification

Bacteria were isolated from two biological niches, larval rearing water and bacterial biofilm at the bottom of the rearing pan. The rearing water was 10-fold serially diluted with sterile 1X Phosphate buffered saline (PBS). The biofilm was gently rinsed five times with 1X PBS then the biofilm was scraped with sterile plastic scraper, resuspended with 20 mL of sterile PBS then 10-fold serially diluted. The serial dilutions were then plated on Luria-Bertani (LB) and incubated at room temperature for 2 days. Bacterial colonies with distinct morphologies from each ecological niche were then streaked on LB agar plate. A single colony from each bacterial isolate was then propagated and used for molecular identification using 16s rRNA gene. Briefly, a colony of each isolate was resuspended in 50 µL of sterile water then 1 µL of resuspended bacteria was used in a PCR reaction using Phusion^®^ High-Fidelity DNA Polymerase (New England Biolabs) with forward (16s-8F: 5’ AGAGTTTGATCCTGGCTCAG 3’) and reverse (16s-1492R: 5’ CGGTTACCTTGTTACGACTT3’) primers (Eden et al., 1991). PCR amplification was performed with an initial denaturation of 2 minutes 30 seconds at 98°C, followed with 30 cycles of denaturation at 98°C for 10 seconds, annealing at 52°C for 15 seconds and an extension at 72°C for 1 minute. The amplicons were then purified using QIAquick PCR purification kit then submitted for Sanger DNA sequencing. The obtained bacterial 16s rRNA gene sequences from the forward and reverse primers were manually trimmed and assembled. The assembled sequences were then searched on the NCBI’s Basic Local Alignment Search Tool (BLAST, https://blast.ncbi.nlm.nih.gov/Blast.cgi) (Altschul et al., 1990).

### 2.4 Bacteria culture and bacterial spent media preparation

All isolated bacteria were cultured in LB medium under specific conditions as mentioned for each experiment. To start a culture, a colony of each bacterium was inoculated in 10 mL of LB medium then cultured at 25°C, 200 rpm overnight. The absorbance at 600 nm (OD_600_) was measured for the overnight culture, then 1 mL of culture at OD_600_ = 1 was inoculated into 5 mL of fresh LB medium. For static culture, 18 mL of inoculated culture was added to a 90 mm diameter petri dish then incubated without shaking at specified temperature. For planktonic culture, 18 mL of inoculated culture was added to a 50 mL conical tube then agitated at 200 rpm at specified temperature. After incubation, bacterial cells were removed from the spent media by centrifugation at 4,415 × *g* 4 ºC for 45 minutes. Supernatants were then filtered through 0.22 µm syringe filter (Sartorius). The sterile spent media were then kept at 4 ºC until further use.

### 2.5 Larvicidal activity assay

Second instar larvae of *An. dirus* were filtered and rinsed twice with 250 mL sterile distilled water. Larvicidal assays were performed with 20 mosquitoes in 4 mL of total water volume per treatment per replicate in each well of 6-well plate. Filtered bacteria spent media were added to each well at 10% v/v and mortality was recorded for 72 hours. For the group with antibiotics treatment, Penicillin/Streptomycin solution (Gibco) were added to the well at final concentration of 100 U/mL and 100 µg/mL, respectively.

For *Pseudomonas* sp. challenge, the bacterium was cultured overnight at 25 ºC then the cells were collected by centrifugation at 1500 × *g*, 4 ºC for 15 minutes. Bacterial cells were then washed twice with 1X PBS, diluted then added to larvicidal plate at concentrations as specified in the results section.

### 2.6 Characterization of physical property of secreted larvicidal molecules and preparation of concentrated macromolecule fraction

We first evaluated whether the secreted larvicidal molecules were proteins (macromolecules) or small molecules because they would require different methods of identification. Thus, the larvicidal molecules were characterized based on the heat stability and molecular weight.

The heat stability of the secreted larvicidal molecules was determined by heating the spent media at 45, 70, and 95 ºC for one hour. Heated spent media were then used to test for remaining larvicidal activity.

Proteins (macromolecules) and small molecules were separated using Amicon^®^ Ultra-4 10K Centrifugal Filter Device following the manufacturer’s instructions. Briefly, 4 ml of spent media were added to the filter unit then centrifuged at 1500 × *g*, 4 ºC for 30 minutes. The flow through was collected as small molecule fraction while the molecules retained on the filter device were used as macromolecule fraction. The concentrated macromolecule fraction was reconstituted with either LB or 1X PBS to concentrations as indicated in each experiment.

### 2.7 Cytotoxicity in C6/36 insect cell line

Cytotoxicity of *S. marcescens* spent medium in insect cell line was determined by sulforhodamine B (SRB) assay according to previously published protocol (Vichai and Kirtikara, 2006) with slight modification for C6/36 cell line. The SRB assay measured cellular protein content to determine cell survival after spent medium treatment. Briefly, C6/36 cells were seeded into each well of 96-well plate at 30,000 cells/well in 100 µl of 10% (v/v) *S. marcescens* spent medium in complete L15 medium (L15 supplemented with 10% Fetal bovine serum, 1% MEM non-essential amino acids, 2% Tryptose phosphate broth, 100 U/mL Penicillin and 100 µg/mL Streptomycin) and incubated at 28 ºC for 72 hours. After incubation, cell monolayers were fixed with 10% (w/v) trichloroacetic acid for 45 minutes at 4 ºC and rinsed 4 times with tap water. Fixed cell monolayer was then stained with 0.057% (w/v) SRB in 1% acetic acid for 30 minutes at room temperature in the dark. Stained cell monolayers were then washed with 100 µl 1% acetic acid for four times to remove unbound dye. To solubilize protein-bound SRB, 200 µl of 10 mM Tris (pH 10.5) was added to each well then the plate was shaken for 30 minutes at room temperature. Absorbance at 510 nm was then measured in a microplate reader to determine cell viability. LB was used as an untreated control and 5 µM cycloheximide was used as a positive control of the cytotoxicity assay.

### 2.8 In-gel digestion of secreted S. marcescens macromolecules for LC-MS/MS analysis

The cell-free macromolecules of *S. marcescens* static culture at 25 and 37 ºC were mixed with Laemmli buffer, denatured at 95 ºC for 10 minutes then subjected to SDS-PAGE. After the SDS-PAGE fractionation, protein bands present in the 25 but not in 37 ºC culture condition were excised from the gel for in-gel tryptic digestion followed by LC-MS/MS analyses (Shevchenko et al., 2007). Briefly, the gel bands were excised and dehydrated in 100% acetonitrile. Proteins were reduced by 10 mM dithiothreitol at 56°C for 30 min and alkylated by 20 mM chloroacetamide at room temperature for 20 min. The gel bands were further destained using 50 mM NH_4_HCO_3_ in 50% acetonitrile for 30 min, followed by dehydration in neat acetonitrile. Digestion was carried out by adding 13 ng/µL trypsin in 50 mM NH_4_HCO_3_ to cover the gel pieces, and the reaction was incubated at 37°C overnight. Extraction of cleaved peptides was performed by adding extraction buffer (70% acetonitrile/1% formic acid) to the digests at the ratio of 2:1 (v/v). Then the extracts were lyophilized and redissolved in 0.1% formic acid ready for LC-MS/MS analyses.

### 2.9 LC-MS/MS analyses of secreted S. marcescens macromolecules

The digested peptides (200 ng) were analyzed by an EASY-nLC 1000 HPLC system (Thermo Fisher Scientific) interfaced with a Q Exactive™ Hybrid Quadrupole-Orbitrap mass spectrometer (Thermo Fisher Scientific) via an EASY-Spray ion source (Thermo Fisher Scientific). The separation was performed for 60 min at the flow rate of 300 nL/min using C18-reverse phase column (EASY-Spray™, Thermo Fisher Scientific). The stepwise gradient was set from 5% solvent B (0.1% formic acid in acetonitrile)/ 95% solvent A (0.1% formic acid in H2O) to 20% solvent B/ 80% solvent A for 43 min and then to 40% solvent B within the next 10 min, followed by holding at 95% for 55 - 60 min. Data-dependent acquisition was performed wherein the ten most abundant precursor ions were selected for fragmentation. The precursor masses were scanned by the Orbitrap mass analyzer at the mass range of 400 – 1600 m/z and the resolution of 70,000. Higher-energy collisional dissociation was employed for fragmentation with an exclusion of +1 charged ions. All data were acquired using Thermo Xcalibur 2.2 software.

Mass spectra were searched against NCBI database of *Serratia* sp. (August, 2018) using Proteome Discoverer 2.2 connected to SEQUEST search engine (Eng et al., 1994). Precursor mass tolerance was set to 10 ppm and the product ion mass tolerance was 0.02 Dalton. Trypsin was specified with maximum 2 missed cleavages. Carbamidomethylation was chosen as a fixed modification; while oxidation of methionine was allowed as variable modification. Decoy search was performed using the false discovery rate of 0.01 as a threshold.

### 2.10 Chitinase activity assay

Chitinase secretion of *S. marcescens* when cultured in different culture temperature was demonstrated by the ability of the bacterium to generate clear zone on LB agar supplemented with 0.5% chitin. *S. marcescens* inoculated plates were cultured at 25°C or 37°C for 10 days before visualization of clear zone.

Chitinase activity of spent medium was determined by measuring the ability of the spent medium to digest chitin into a reducing sugar, *N*-acetyl-*D*-glucosamine. To set up the chitinase activity assay, 100 µL of cell-free spent culture media served as a crude enzyme source was mixed with 100 µL of 1% (w/v) chitin powder (Sigma, USA) in 50 mM sodium phosphate buffer (pH 6.5). The reaction mixture was incubated at 30ºC for 90 min. The amount of reducing sugar was then determined by using *p*-hydroxybenzoic acid hydrazide (pHBAH) assay (Lever, 1972). A stock solution of 5% (w/v) pHBAH (Sigma, USA) was prepared in acid medium (0.5 M HCl) and diluted at the moment of analysis to 1% (w/v) in alkaline medium (0.5 M NaOH). Samples were mixed with 1% (w/v) pHBAH in alkaline solution (0.5 M NaOH) at a ratio of 1:3 and the mixture was heated in the boiling bath for 5 min to develop the color. After cooling, the solution was centrifuged at 13,800 × *g* for 2 min. The absorbance at 410 nm of the supernatant was then measured using spectrophotometer. *N*-acetyl-*D*-glucosamine standard solution was prepared in the same manner at concentrations of 0, 0.2, 0.4, 0.6, 0.8, and 1 mg/ml in distilled water. One unit of enzyme activity was defined as the amount of enzyme which releases 1 µg of reducing sugar per hour under the assay conditions.

### 2.11 Statistical analyses

Statistical significance of the survival analyses was calculated using Log-rank test followed by a Bonferroni post-hoc analysis using ‘survminer ‘ package (Kassambara et al., 2019) in R (R Core Team, 2020).

Concentration of *Pseudomonas* sp. treatment resulting in 50% larval mortality (LC_50_) and 90% larval mortality (LC_90_) was calculated by Probit analysis with Abbot’s modification using ‘ecotox’ package (Scholefield et al., 2008) in R (R Core Team, 2020).

Statistical analysis comparing chitinase activity level and inhibition of chitinase activity was performed using One-way ANOVA with Tukey multiple comparison test using multcompView (Graves and Dorai-Raj, 2019) package in R (R Core Team, 2020).

## 3. Results

### 3.1 Isolation of bacteria from An. dirus larval rearing pan

A total of seven bacteria species were isolated and identified from the rearing pan by 16s rRNA gene sequencing. These include one Actinobacteria: *Leucobacter* spp. (Leu_spp), two Bactereoidetes: *Sphingobacterium* spp. (Sph_spp) and *Sphingobacterium multivorum* (Sch_m); and four proteobacteria: *Acinetobacter soli* (Aci_s), *Ochrobactrum* spp. (Och_spp), *Serratia marcescens* (Ser_m), and *Stenotrophomonas maltophilia* (Ste_m) as summarized in Table 1.

**Table 1.**
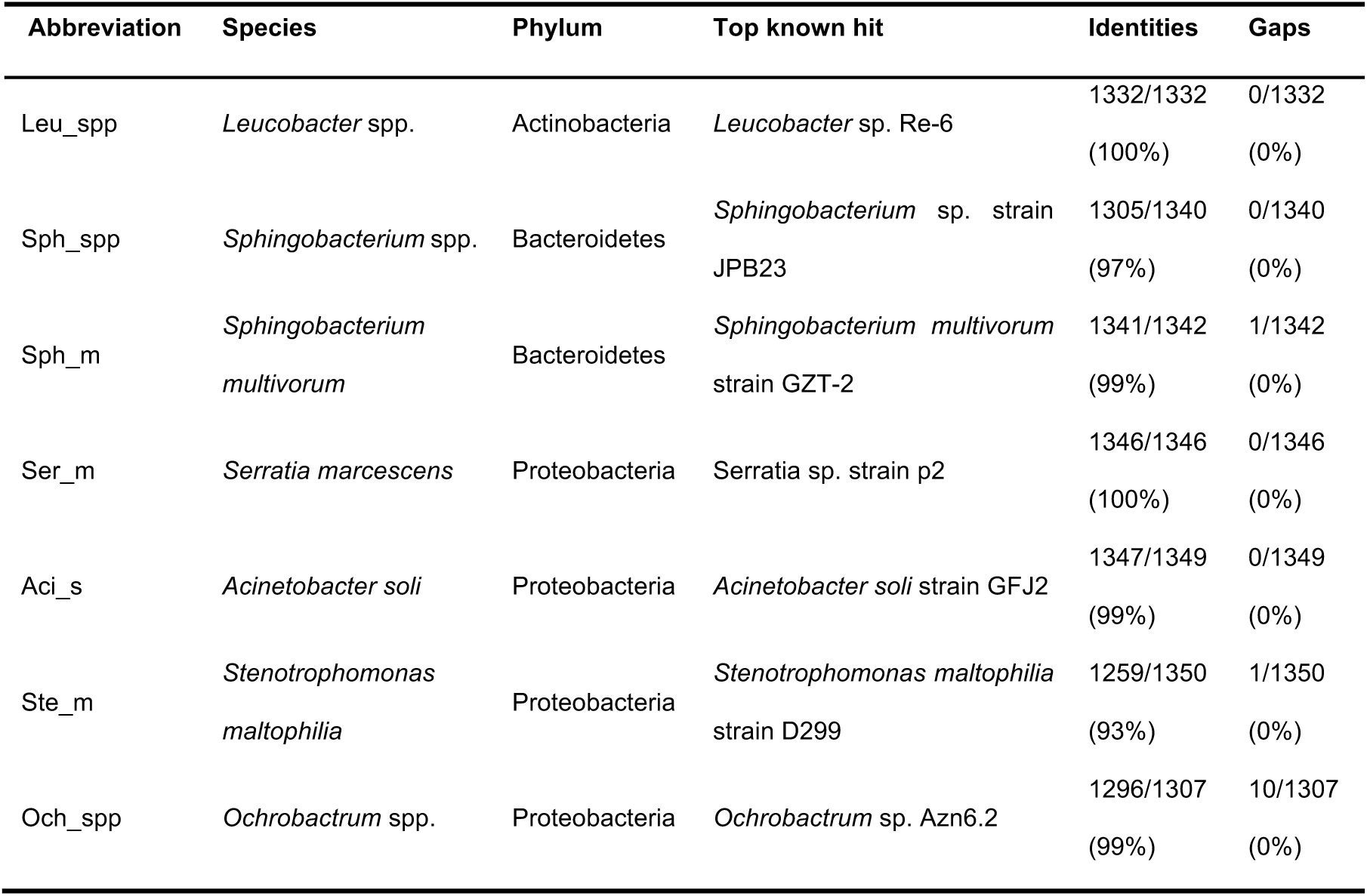
Bacterial isolates from *An. dirus* larval rearing pan identified by 16s rRNA gene sequencing

| Abbreviation | Species | Phylum | Top known hit | Identities | Gaps |
| --- | --- | --- | --- | --- | --- |
| Leu_spp | *Leucobacter* spp. | Actinobacteria | *Leucobacter* sp. Re-6 | 1332/1332<br>(100%) | 0/1332<br>(0%) |
| Sph_spp | *Sphingobacterium* spp. | Bacteroidetes | *Sphingobacterium* sp. strain JPB23 | 1305/1340<br>(97%) | 0/1340<br>(0%) |
| Sph_m | *Sphingobacterium multivorum* | Bacteroidetes | *Sphingobacterium multivorum* strain GZT-2 | 1341/1342<br>(99%) | 1/1342<br>(0%) |
| Ser_m | *Serratia marcescens* | Proteobacteria | *Serratia* sp. strain p2 | 1346/1346<br>(100%) | 0/1346<br>(0%) |
| Aci_s | *Acinetobacter soli* | Proteobacteria | *Acinetobacter soli* strain GFJ2 | 1347/1349<br>(99%) | 0/1349<br>(0%) |
| Ste_m | *Stenotrophomonas maltophilia* | Proteobacteria | *Stenotrophomonas maltophilia* strain D299 | 1259/1350<br>(93%) | 1/1350<br>(0%) |
| Och_spp | *Ochrobactrum* spp. | Proteobacteria | *Ochrobactrum* sp. Azn6.2 | 1296/1307<br>(99%) | 10/1307<br>(0%) |

### 3.2 S. marcescens secreted larvicidal molecule into spent medium when cultured under static condition

Because larval mortality was observed a few days after bacterial bloom, we hypothesized that the isolated bacteria could secrete larvicidal molecule(s) after the bacteria grow to a certain density; at a time when biofilm was observed at the bottom of the pan for a few days. Therefore, the isolated bacteria were cultured under either static or planktonic conditions for three days before spent media collection for the screening of larvicidal activity.

None of the cell-free spent media obtained from planktonic condition had any toxicity against *An. dirus* larvae (Figure 1A). However, for the stationary culture, only the cell-free spent media from *S. marcescens* had a detrimental effect on the survival of the larvae (Figure 1A). Next, spent medium of *S. marcescens* cultured under planktonic and static conditions was collected over-time after 1, 2, or 3 days of incubation. Again, the larvicidal molecule was secreted into the *S. marcescens* only under static culture condition and the larvicidal activity of the spent medium increased over time starting from the second day after inoculation (Figure 1B).

**Figure 1.**
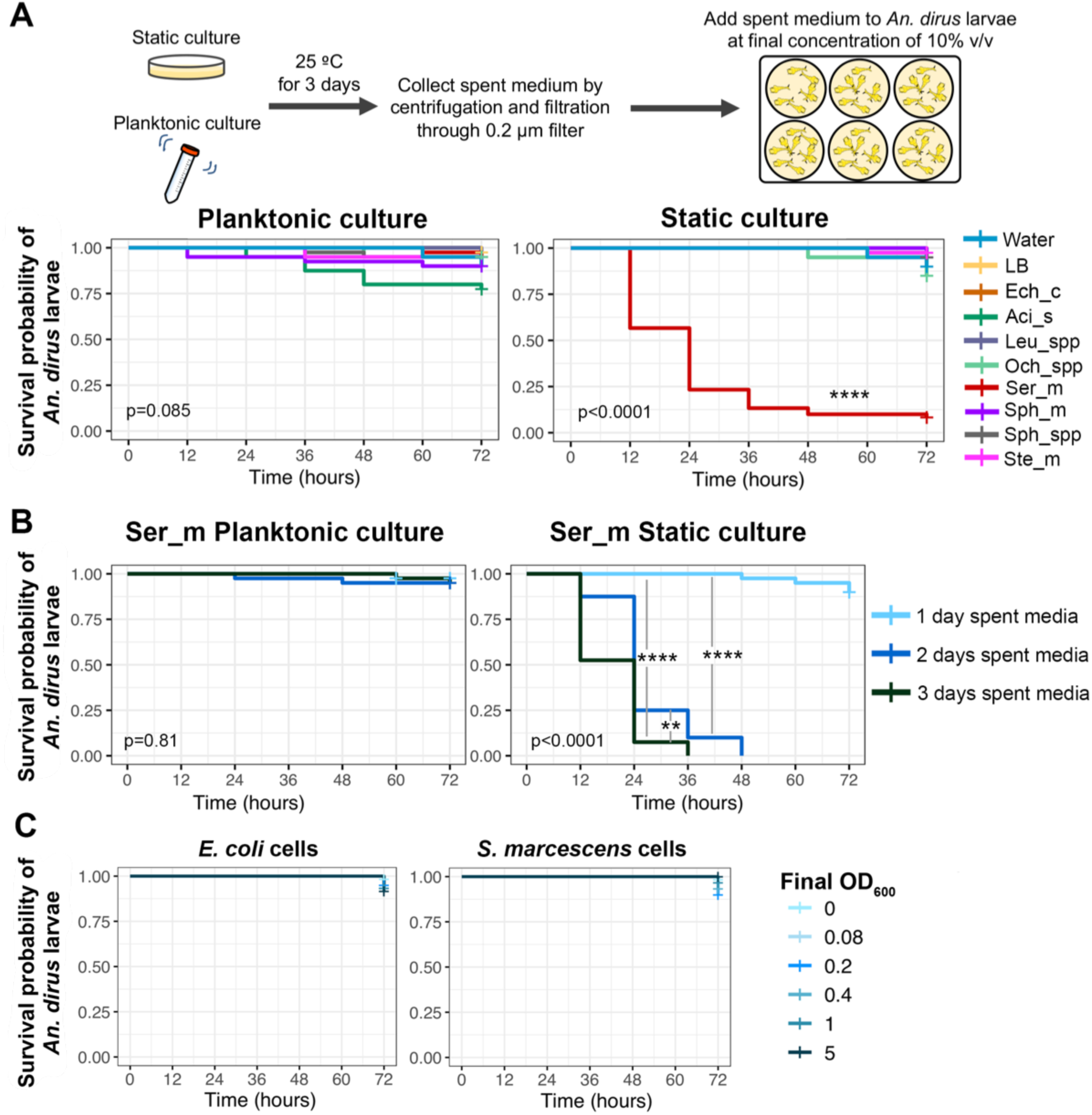
Survival analyses of *An. dirus* mosquito larvae after treatment with bacterial spent media at 10% (v/v) concentration. (A) Survival curve of *An. dirus* larvae after treatment with bacterial spent media cultured under planktonic condition or static condition. The spent media were collected at three days after incubation under planktonic or static state at 25 ºC Ech_c: *Escherichia coli* strain XL10, Aci_s: *Acinetobactor soli*, Leu_spp: Leucobactor spp., Och_spp: Ochrobatrum spp., Ser_m: *Serratia marcescens*, Sph_m: *Sphingobacterium multivorum*, Sph_spp: *Sphingobacterium* spp., Ste_m: *Stenotrophomonas maltophilia*. LB medium or water was used as a control at final concentration of 10% (v/v). The graphs represent data from three independent biological replicates (B) Survival curve of *An. dirus* larvae after treatment with *S. marcescens* spent medium cultured under planktonic or static condition and collected after 1, 2, or 3 days after inoculation. The graphs represent data from two independent biological replicates. (C) Survival curve of *An. dirus* larvae after treatment with *E. coli* or *S. marcescens* cells at various concentration. Bacterial cells were cultured under static condition at 25 ºC then washed twice with PBS. The graphs represent data from three independent biological replicates. Statistical significance of the survival analyses was calculated using Log-rank test followed by a Bonferroni post-hoc analysis using ‘survminer ‘package in R. Significance level: **: p<0.01, ****: p<0.0001.

To investigate if the *S. marcescens* cells are entomopathogenic to *An. dirus* larvae, the bacterium was cultured under a static condition then washed with 1X PBS. Washed bacterial cells were then added to larval culture at various concentrations (Figure 1C). Even at OD_600_ of 5, *S. marcescens* cells did not cause significant mortality to *An. dirus* larvae. The *Escherichia coli* cells used as a control also did not cause larval mortality. These results suggested that *S. marcescens* only cause larval mortality through the secretion of entomopathogenic molecules.

### 3.3 Larvicidal molecules from S. marcescens are heat sensitive macromolecules

Because the nature of larvicidal molecule would dictate subsequent methods for identification and characterization, the larvicidal molecule was then next broadly characterized based on heat sensitivity and molecular weight. The *S. marcescens* filtered spent medium was heated at 45, 70 and 95 ºC for one hour then used to determine larvicidal activity at 10% (v/v) (Figure 2A). The larvicidal molecules secreted by *S. marcescens* are heat-sensitive as the heat treatment at 45 ºC for 1 hour significantly reduced the 3-day mortality rate of treated larvae from 100% to 35%. Heat treatment at 70 and 95 ºC completely destroyed the larvicidal activity. To further characterize if the heat-labile active larvicidal elements were small or large macromolecules, we fractionated the active spent media by size exclusion using Amicon^®^ Ultra-4 10K filter device and tested their larvicidal activity. The result shows that only the macromolecule fraction with molecular weight larger than 10 kDa possesses larvicidal activity (Figure 2B).

**Figure 2.**
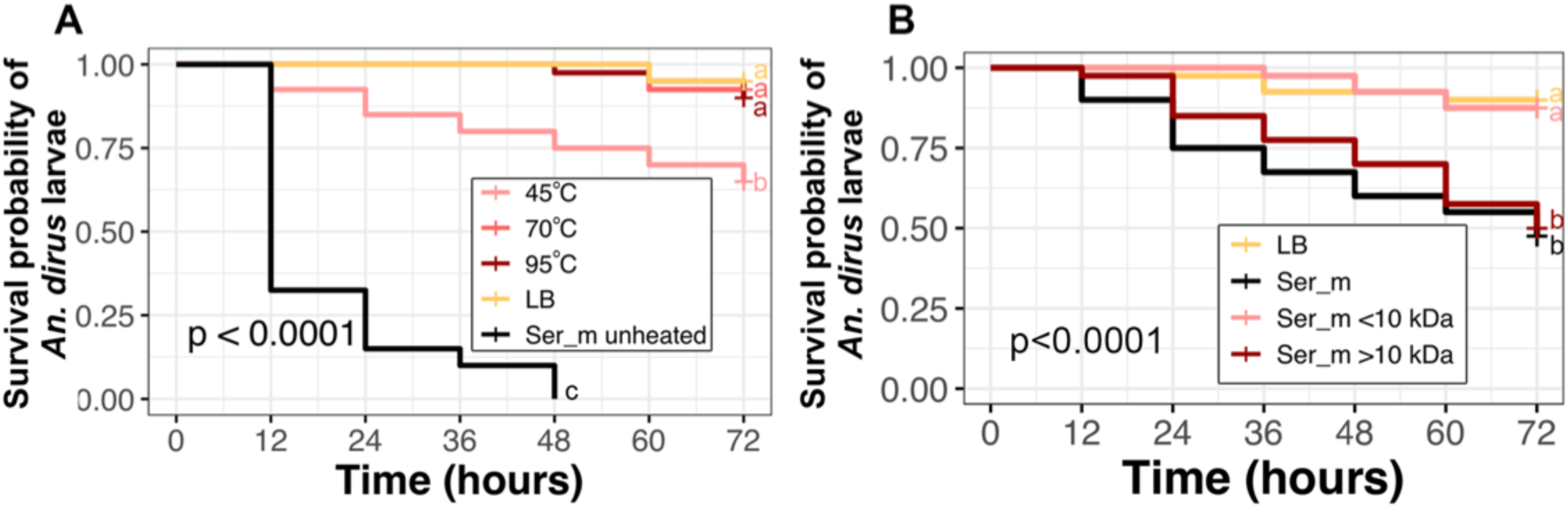
Larvicidal molecule(s) secreted by *S. marcescens* are heat sensitive macromolecule(s). (A) Heat sensitivity of larvicidal activity from *S. marcescens* spent medium. The *S. marcescens* spent medium was heated at 45, 70, and 95 ºC for one hour before testing for larvicidal activity at 10% (v/v) final concentration. LB medium was used as a control at a final concentration of 10% (v/v). The graphs represent data from two independent biological replicates (B) Larvicidal activity of *S. marcescens* spent medium after separation of small (<10 kDa) and macromolecules (>10 kDa) using Amicon^®^ Ultra-4 10K filter device at 10% (v/v). LB medium was used as a control at final concentration of 10% (v/v). The graphs represent data from two independent biological replicates. Statistical significance of the survival analyses was calculated using Log-rank test followed by a Bonferroni post-hoc analysis using ‘survminer ‘package in R.

### 3.4 Larvicidal activity of S. marcescens spent medium is not likely to be caused by cytotoxicity

To test the cytotoxicity of *S. marcescens* spent medium against insect cells, the spent medium was used to treat C6/36 cells at the final concentration of 10% (v/v) spent medium in complete L15 medium at 28 ºC for 72 hours. When compared to LB medium treatment, the cell viability of C6/36 treated with *E. coli* spent medium, *S. marcescens* spent medium, and cycloheximide were 111.25±8.23%, 76.79±15.02%, and 7.82±2.06%, respectively (Figure 3). Although statistically significant, the low cytotoxicity level of *S. marcescens* spent medium against C6/36 cells could not support the strong larvicidal activity as shown in our experiments. This could be because the cell does not carry the essential molecule/pathway targeted by *S. marcescens*’ entomopathogenic molecules. Another possible explanation is that the death of *An. dirus* larvae is due to damage at the physiological level rather than direct toxicity to insect cells.

**Figure 3.**
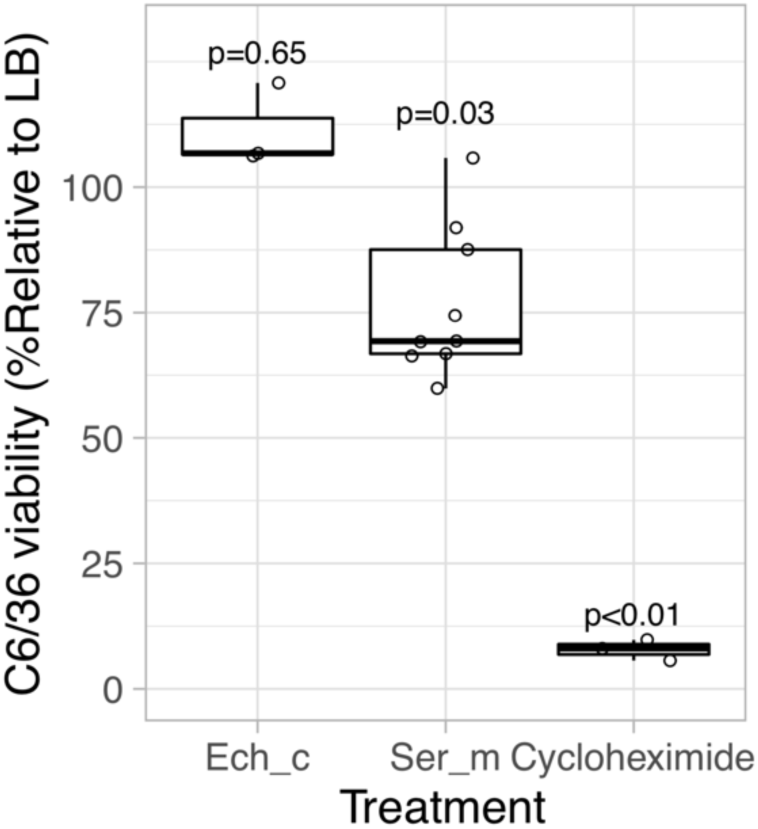
C6/36 cell viability assay of *E. coli* spent medium (Ech_c), *S. marcescens* spent medium (Ser_m), and 5 µM cycloheximide. The cell viability was shown relative to LB medium treatment. Spent medium and LB were used at 10% (v/v) final concentration. P-values were calculated using raw data from sulforhodamine B assay by One-way ANOVA with TukeyHSD post hoc test with multcompView package in R software.

### 3.5 S. marcescens spent medium can directly kill An. dirus larvae and also enhance larvae’s susceptibility to bacterial infection

In previous experiments, LB medium was used as a control in the larvicidal assays. We observed that the presence of LB or filtered spent medium allowed bacteria in the environment to grow in these treatments (Figure 1 and 2). Although these bacteria did not cause larval death in the LB-treated group, they might interfere with the larvicidal activity caused by *S. marcescens* spent medium treatment.

To test whether these naturally growing bacteria have any impact on larvicidal activity by *S. marcescens*, penicillin (100 U/mL) and streptomycin (100 µg/mL) were added in the larvicidal assay experiment to prevent bacterial growth. The addition of antibiotics significantly inhibited bacterial contamination because water in each well of the larvicidal assay remained clear throughout the experiment. In the absence of natural bacterial growth, the macromolecules secreted by *S. marcescens* can still kill *An. dirus* larvae confirming its intrinsic toxicity (Figure 4A). Interestingly, the observed mortality in the antibiotics-treated group was significantly lower than the group without antibiotics treatment. In addition to antibiotics treatment, resuspension of *S. marcescens* macromolecules in PBS instead of LB also reduced natural bacterial growth in larvicidal assays. Even without obvious natural bacterial growth, *S. marcescens* macromolecule resuspended in PBS could kill *An. dirus* larvae in a concentration-dependent manner (Figure 4B). These results suggest that the larvicidal molecule has intrinsic larvicidal activity, and also increases larvae’s susceptibility to bacterial infection.

**Figure 4.**
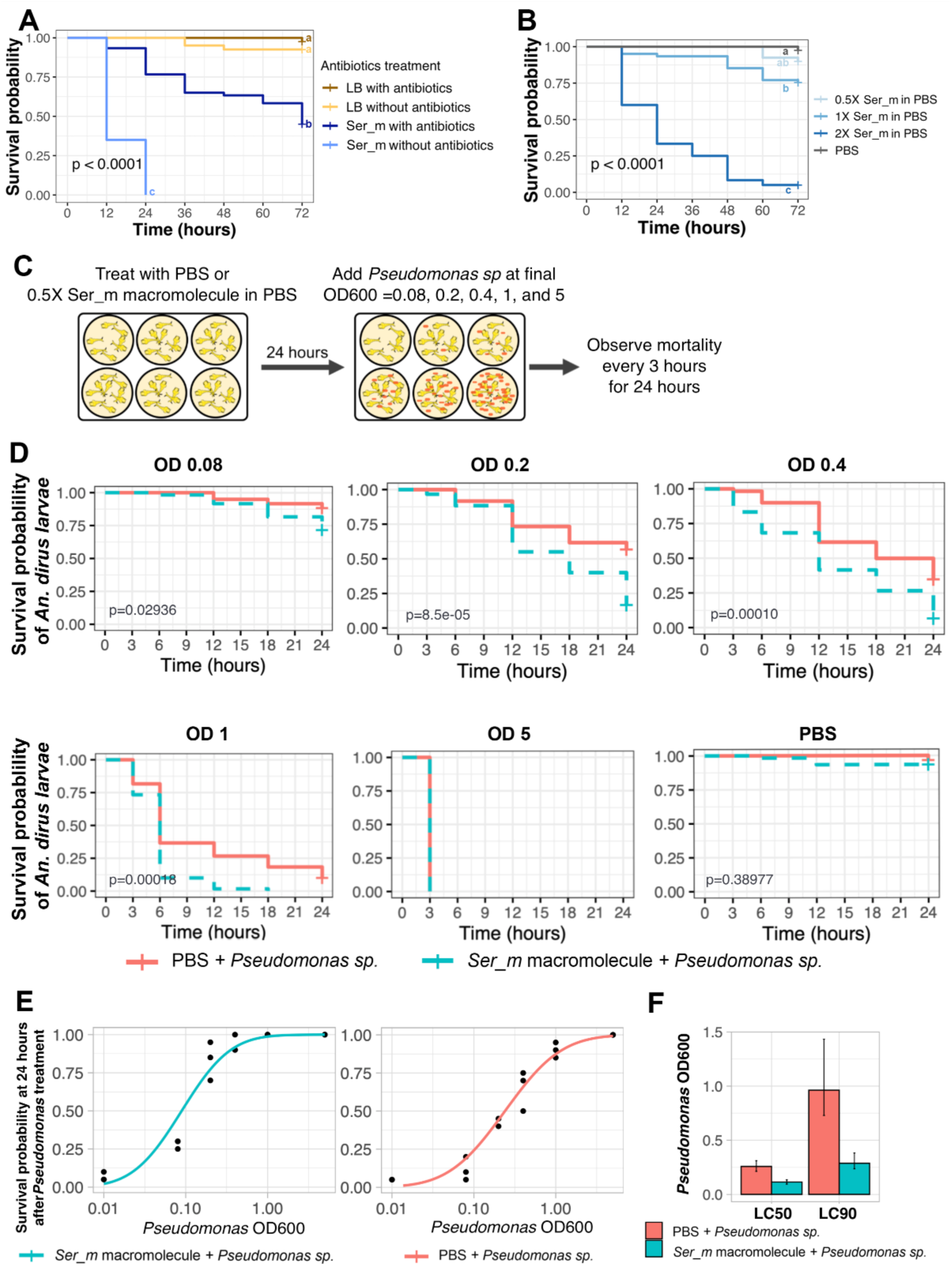
Intrinsic larvicidal activity of *S. marcescens* macromolecule and its enhancing effect on larvae’s susceptibility to bacterial infection. (A) Survival of *An. dirus* larvae treated with 2X concentration of macromolecules from *S. marcescens* spent medium diluted in LB with or without antibiotics. LB with or without antibiotics were used as a control. The graphs represent data from two independent biological replicates (B) *An. dirus* larvae treated with *S. marcescens* macromolecules reconstituted to the final concentration of 0.5X, 1X, and 2X in PBS. The graphs represent data from two independent biological replicates. (C) Schematic diagram of the treatment of *An. dirus* larvae with sublethal concentration of *S. marcescens* secreted macromolecule followed by *Pseudomonas* sp. treatment. The larvae were treated for 24 hours with 0.5x concentration of macromolecules from *S. marcescens* diluted in PBS buffer before addition of *Pseudomonas* sp. at final concentration of OD_600_ = 0.08, 0.2, 0.4, 1, and 5. Mortality of the larvae was monitored every 3 hours for 24 hours. The graphs represent data from three independent biological replicates. (D) Kaplan-Meier curve of *An. dirus* larvae from *Pseudomonas* sp. after treatment with sublethal concentration of *S. marcescens* macromolecules comparing to the PBS control. (E) Probit analysis of *An. dirus* survival probability at 24 hours after treatment of *S. marcescens* macromolecules. (F) LC_50_ and LC_90_ of *Pseudomonas* sp. treatment in *An. dirus* calculated from Probit analysis.

To confirm the hypothesis that *S. marcescens* secreted macromolecules increases larvae’s susceptibility to bacterial infection, we pretreated *An. dirus* larvae with a sublethal concentration of *S. marcescens* secreted macromolecule (0.5X diluted in PBS; see Figure 4B) for 24 hours then added an entomopathogenic strain of *Pseudomonas* sp. at the final bacterial density of OD_600_ = 5, 1, 0.4, 0.2, and 0.08 (Figure 4C). *An. dirus* larvae pretreated with *S. marcescens* secreted macromolecules significantly increased the mortality rate caused by *Pseudomonas* sp. in all treatment except at OD 5, which the entomopathogenicity was too high to observe the difference (Figure 4D). The *Pseudomonas* sp. LC_50_ of the *S. marcescens* macromolecule-treated group was 0.115 OD_600_, a 55% decrease from 0.258 OD_600_ in the PBS-treated group. Similarly, the *Pseudomonas* sp. LC_90_ of the *S. marcescens* macromolecule-treated group was 0.287 OD_600_, a 70% decrease from 0.962 OD_600_ in the PBS-treated group (Figure 4E and 4F). These results confirm significant effect of *S. marcescens* macromolecules on susceptibility of *An. dirus* larvae to bacterial infection, even at sublethal concentration.

### 3.6 *S. marcescens* secreted different sets of macromolecules when cultured under different conditions

Because we observed different levels of larvicidal activity from *S. marcescens* spent medium when cultured under different conditions, the comparison of macromolecules from these culture conditions would help us to pinpoint macromolecule(s) that has larvicidal activity. Although the planktonic condition did not induce secretion of a larvicidal molecule, *S. marcescens* barely secreted proteins under planktonic condition. Further comparison between these two conditions would yield several protein candidates for further studies.

To identify other culture conditions that might result in differential larvicidal activity, *S. marcescens* was cultured under static condition at different temperatures, 25 ºC and 37 ºC. Here, *S. marcescens* spent medium from 25 ºC incubation could kill *An. dirus* larvae while those cultured at 37 ºC could not (Figure 5A).

**Figure 5.**
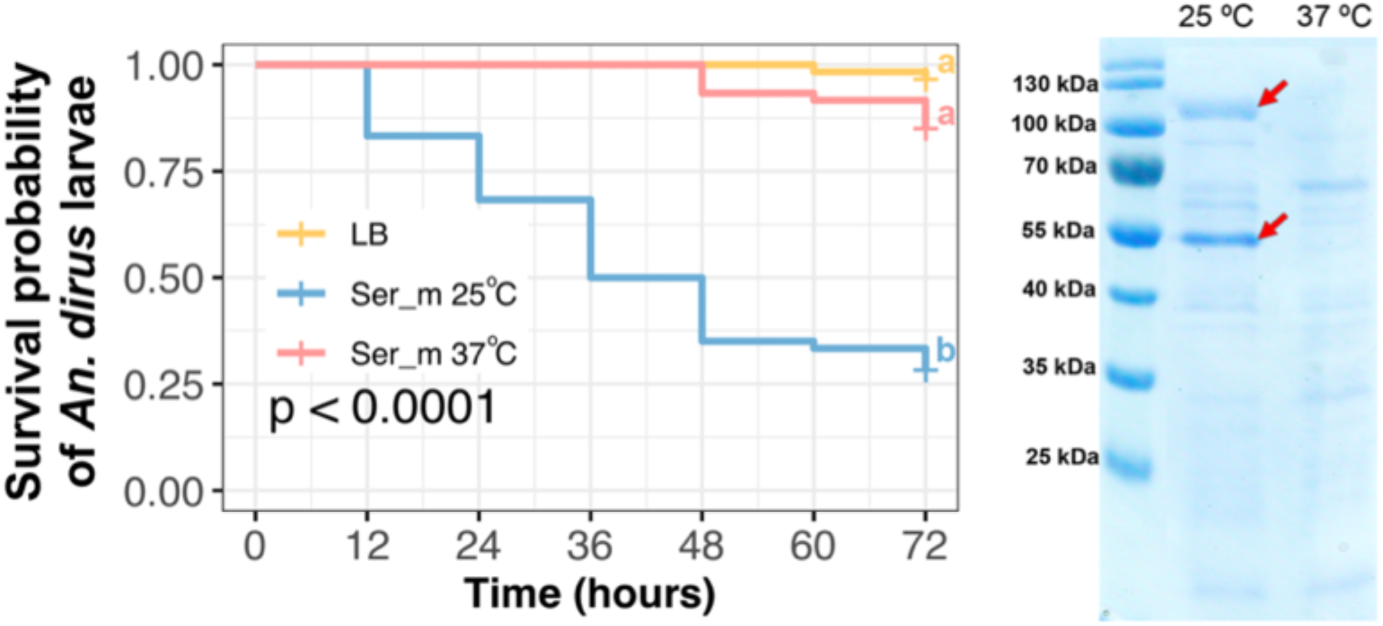
*S. marcescens* secreted larvicidal molecules when cultured under static condition at 25 ºC but not 37 ºC. (A) Survival curve of *An. dirus* larvae after treatment with *S. marcescens* spent media cultured under static condition at 25 ºC or 37 ºC. The graphs represent data from three independent biological replicates. Statistical significance of the survival analyses was calculated using Log-rank test followed by a Bonferroni post-hoc analysis using ‘survminer ‘package in R. (B) SDS-PAGE of *S. marcescens* spent media cultured under static condition at 25 ºC or 37 ºC. Two red arrows indicate major protein bands that were expressed when the bacterium was cultured at 25 ºC but not 37 ºC.

Pattern of secreted protein from *S. marcescens* cultured at 25 ºC and 37 ºC was then compared by SDS-PAGE (Figure 5B). Two major protein bands (approximately 110 and 55 kDa) were observed in the spent medium from 25ºC culture but not 37ºC (Figure 5B). These two protein bands were then excised from the gel then subjected to LC-MS/MS for identification.

The 110 kDa protein band consists of several protease enzymes including aminopeptidase N, serralysin, and peptidase M60 viral enhancin protein; chitinase enzymes including chain A chitobiase; cell surface protein; DUF4214 domain-containing protein; hemolysin; DUF4214 domain-containing protein; peptide ABC transporter substrate-binding protein; aconitate hydratase; DNA-directed RNA polymerase; formate dehydrogenase-N subunit alpha; glycine dehydrogenase; translation elongation factor Tu; phosphoenolpyruvate carboxykinase; LysR family transcriptional regulator; short-chain dehydrogenase/reductase SDR; phosphorylcholine phosphatase; acyl-CoA dehydrogenase (Supplementary table 1).

The 55 kDa protein band also consists of proteases including serralysin, proline dipeptidase, extracellular serine protease precursor, secreted alkaline metalloproteinase, and Xaa-Pro aminopeptidase; chitinase enzyme; serine 3-dehydrogenase; DUF4214 domain-containing protein; arginine-tRNA ligase; succinylarginine dihydrolase; cell surface protein; histidine ammonia-lyase; capsule assembly Wzi family protein; glutathione reductase; phosphoenolpyruvate carboxykinase; flagellin; F0F1 ATP synthase subunit beta; and hypothetical protein (Supplementary table 2).

### 3.7 Protease inhibitors reduced larvicidal activity of S. marcescens spent medium

The LC-MS results indicated that *S. marcescens* secreted several proteases when cultured at 25 ºC under a static condition. Some of these proteases have been reported for insecticidal activity in other insects (Baumann, 2013; Wang and Granados, 1997). Therefore, we added the Complete protease inhibitor (Roche) supplemented with 5 mM EDTA in the larvicidal assays to determine whether these proteases contributed to *S. marcescens* larvicidal activity. We found that protease inhibitors significantly reduced the three-day mortality of the larvae to 50% as compared to 87.5% of the control group without protease inhibitors. This suggests that the larvicidal activity of proteases was partially involved in the observed larval mortality (Figure 6).

**Figure 6.**
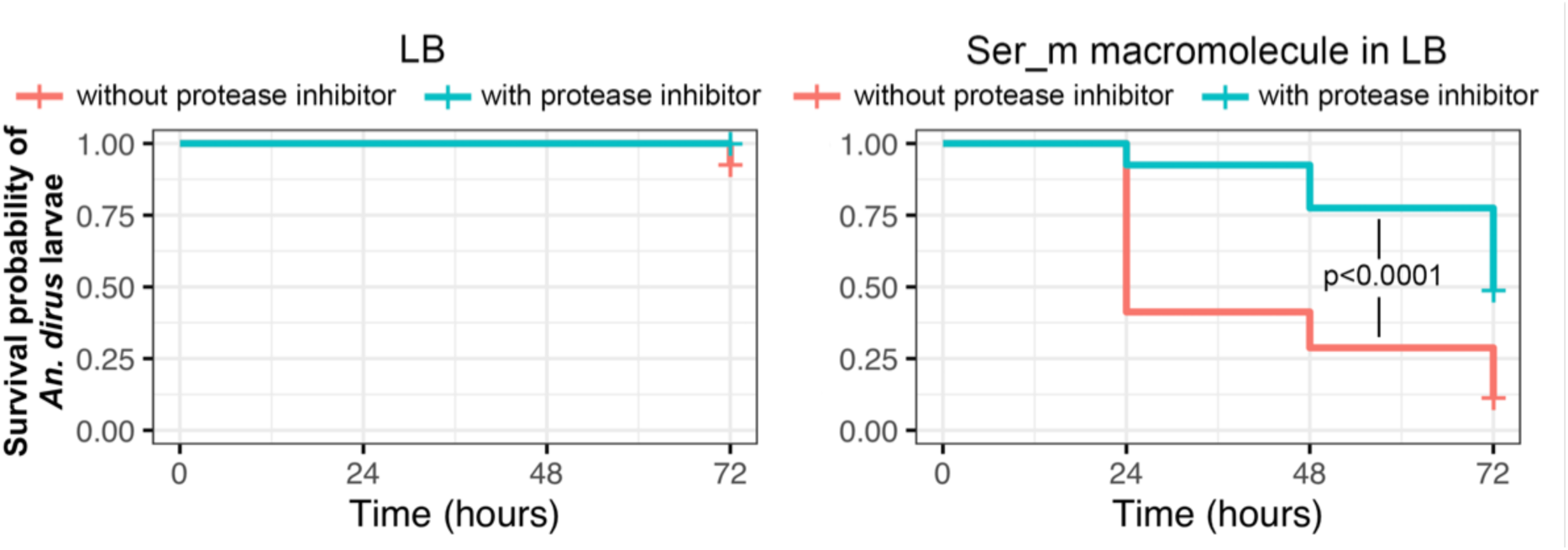
Protease inhibitor reduced larvicidal activity of *S. marcescens* secreted macromolecule. The graphs represent data from four independent biological replicates. Statistical significance of the survival analyses was calculated using Log-rank test followed by a Bonferroni post-hoc analysis using ‘survminer ‘package in R.

### 3.8 Chitinase inhibitor reduced larvicidal activity of S. marcescens spent medium

Because the LC-MS analysis suggested that our strain of *S. marcescens* expressed chitinase and chitobiose enzymes when cultured under static condition at 25 ºC but not 37 ºC, we then inoculated the bacterium on LB agar supplemented with 0.5% chitin then incubated at 25 ºC or 37 ºC to determined chitinase secretion. After ten days of incubation, *S. marcescens* cultured at 25 ºC produced a clear zone while the incubation at 37 ºC did not (Figure 7A). Chitinase activity was also determined in the liquid static culture by pHBAH assay. Chitinase activity of *S. marcescens* spent medium increased overtime during the incubation under static condition at 25 ºC and saturated after 3 days of incubation (Figure 7B). This result coincides well with the increase in larvicidal activity during the first three days of *S. marcescens* incubation (Figure 1B).

**Figure 7.**
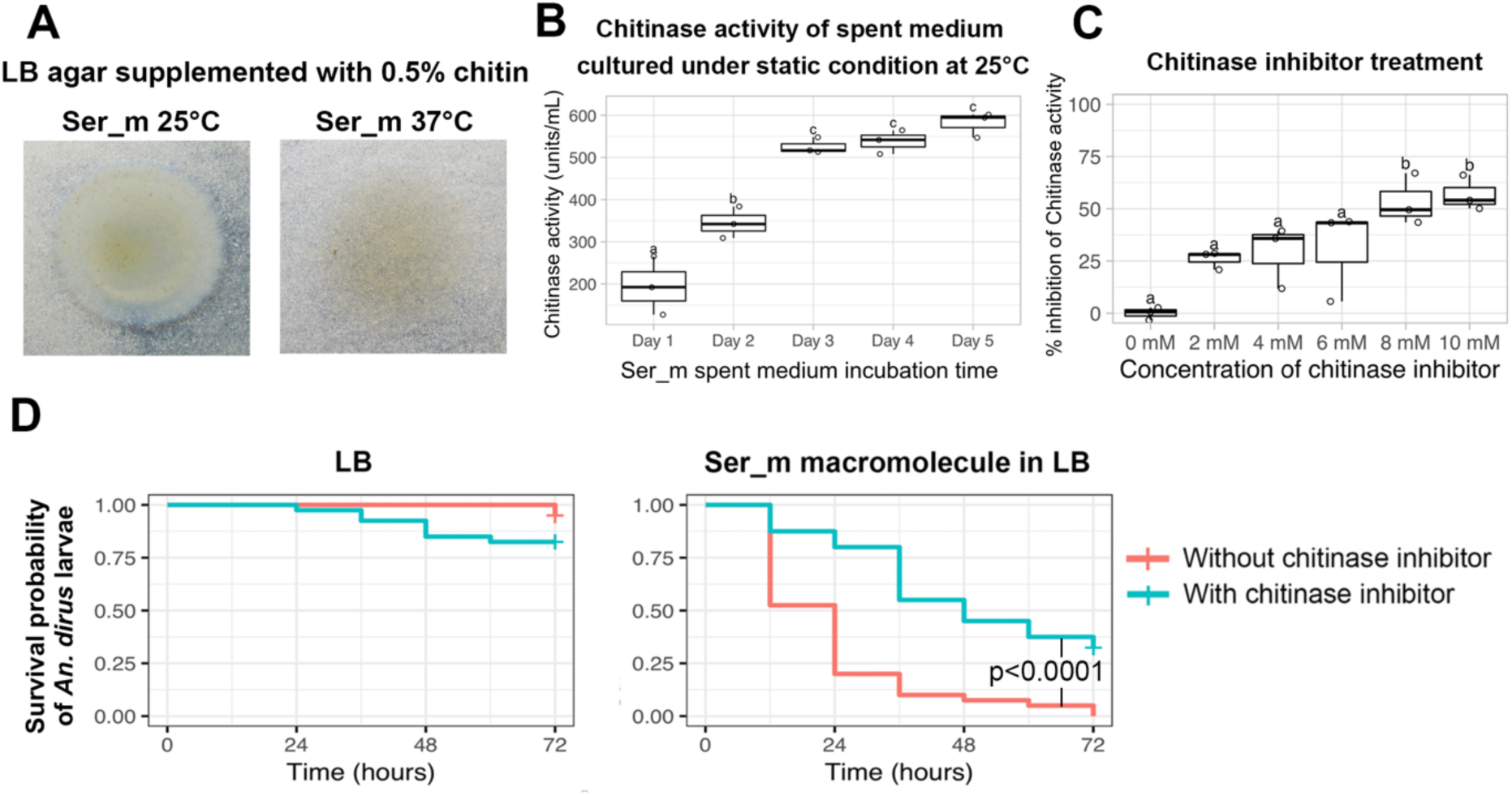
*S. marcescens* secreted macromolecules have chitinase activity and chitinase inhibitor reduced larvicidal activity. (A) *S. marcescens* secreted chitinase that resulted in a clear zone around bacterium colony when cultured at 25 ºC but not 37 ºC. (B) *S. marcescens* spent medium accumulated chitinase activity over time and saturated after three days of incubation. One unit of chitinase activity was defined as the amount of enzyme which releases 1 µg of reducing sugar per hour as measured by pHBAH assay. (C) Cyclo-Gly-Pro inhibit chitinase activity in a concentration-dependent manner. Statistical analyses comparing chitinase activity and inhibition of chitinase activity were performed using One-way ANOVA with Tukey multiple comparison test using multcompView package in R. (D) Chitinase inhibitor reduced larvicidal activity of the *S. marcescens* secreted macromolecule. The graphs represent data from two independent biological replicates. Statistical significance of the survival analyses was calculated using Log-rank test followed by a Bonferroni post-hoc analysis using ‘survminer ‘package in R.

Cyclic dipeptides such as Cyclo-Gly-Pro have previously been reported to inhibit chitinase enzyme activity (Houston et al., 2004). This compound was then used to confirm that the larvicidal activity was due to chitinase secretion. The results in Figure 6C show that Cyclo-Gly-Pro can inhibit chitinase activity of *S. marcescens* spent medium in a concentration-dependent manner. At 8 mM, Cyclo-Gly-Pro inhibited chitinase activity by 53.36±12.22%, which was not significantly different from 56.81±8.34% inhibition by 10 mM Cyclo-Gly-Pro (Figure 7C). Thus, 8 mM Cyclo-Gly-Pro was used in the following larvicidal assays. Larvicidal activity was significantly reduced when 8 mM Cyclo-Gly-Pro was added to the spent medium compared to the group without the chitinase inhibitor, albeit not to the same level as the LB control (Figure 7D). This might be due to the incomplete inhibition of the chitinase activity or other enzymes might be contributing to the larvicidal activity in addition to chitinases.

### 3.9 Combination of protease inhibitors and chitinase inhibitor could not completely inhibit larvicidal activity

Since both the protease inhibitors and chitinase inhibitors could partially reduced larvicidal activity, the combination of both inhibitors rewas used to determine if simultaneous inhibition of proteases and chitinases could completely inhibit the larvicidal activity (Figure 8). The combination of protease and chitinase inhibitors significantly reduced larvae mortality to 33.33% from 93.33% in the no inhibitor control. Although not significantly different, the combination of protease and chitinase inhibitors has a trend to further reduce larvicidal activity compared to the treatment with single class of inhibitors (56.66% in the protease inhibitor treatment and 48.33% in the chitinase inhibitor treatment).

**Figure 8:**
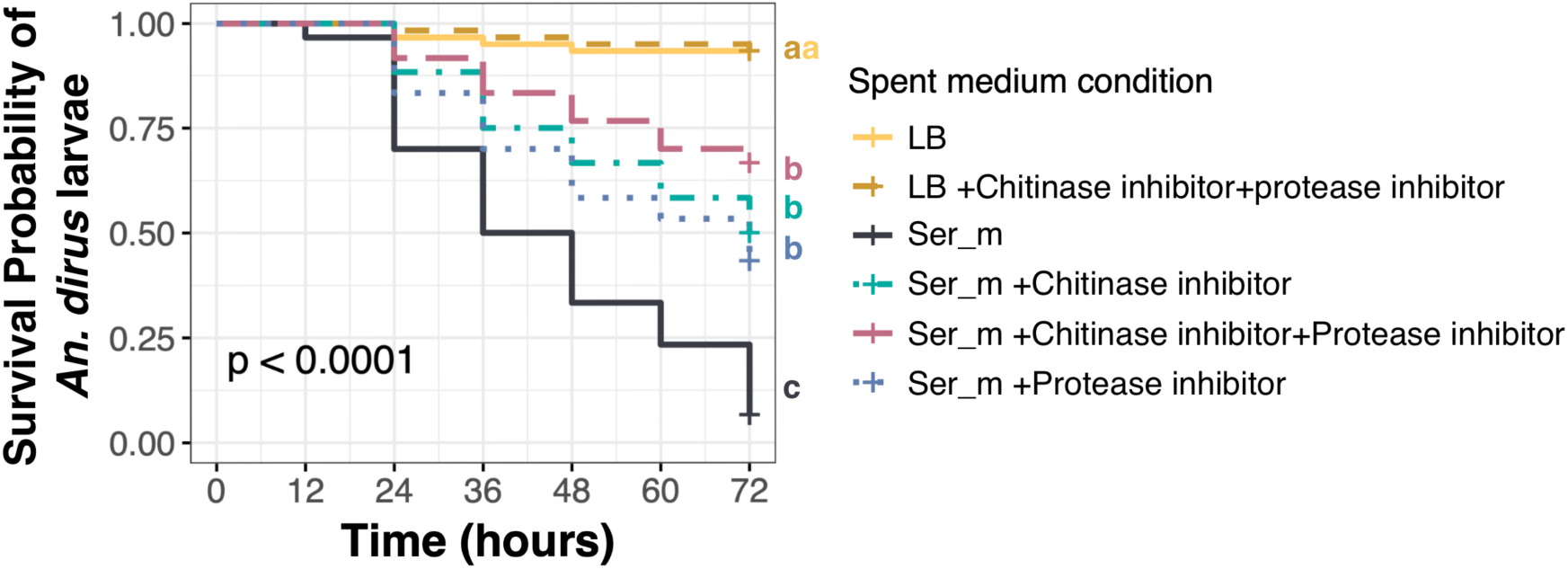
Combination of protease inhibitor and chitinase inhibitor could not completely inhibit larvicidal activity. The graph represents data from three independent biological replicates. Statistical significance of the survival analyses was calculated using Log-rank test followed by a Bonferroni post-hoc analysis using ‘survminer ‘package in R.

## 4. Discussion

In this study, a strain of non-prodigiosin producing *S. marcescens* was identified and found to secrete molecules with larvicidal activity against *Anopheles dirus*, a malaria vector in Southeast Asia. *S. marcescens* is a Gram-negative bacterium commonly found in soil and water but not as a human fecal microbiome (Donnenberg, 2015). With such natural habitats, it is often associated with insects including mosquitoes (Gendrin and Christophides, 2013; Wang et al., 2017, 2011). Interestingly, different strains of *S. marcescens* can have different relationship with *Anopheles* mosquitoes raging from symbiotic (Wang et al., 2017) to entomopathogenic (Bahia et al., 2014). The cells of *S. marcescens* isolated in the present study did not kill *An. dirus* larvae directly, but instead can secrete larvicidal molecules into the culture medium when cultured under static condition for at least 2-3 days. The results were in accordance with our original observations that the larval death occurred a few days after larval water became turbid. The fact that the larvicidal molecules were secreted is an advantage for application purposes, given that the secreted factors can be applied instead of the bacterium. Thus preventing or limiting the risk of nosocomial infections (Golemi-Kotra, 2008). The secreted factors can also be expressed heterologously in other expression systems that are more suited for industrial scale production.

*S. marcescens* has been extensively studied for its potential as insect pest control agent in agriculture due to its insecticidal property (Bidari et al., 2017; Li et al., 2011; Miller et al., 2008; Pineda-Castellanos et al., 2015; Tu et al., 2010). The bacterium was known to secrete many insect virulence factors including several enzymes, as well as secondary metabolite including a red pigment prodigiosin. Prodigiosin is a metabolite possessing several bioactivities including antimalarial and insecticidal/mosquito larvicidal properties (Bidari et al., 2017; Darshan and Manonmani, 2015; Patil et al., 2011; Suryawanshi et al., 2015). The fact that our *S. marcescens* isolate doesn’t produce red pigment suggested that the bacterium secretes other mosquito larvicidal molecules. This finding is in line with a previous study reporting that level of prodigiosin production did not correlate with *S. marcescens* insecticidal activity against silk worm larva (Zhou et al., 2016).

The heat-labile larvicidal macromolecules were secreted only when *S. marcescens* was cultured under static condition at lower temperature of 25 ºC but not 37 ºC nor at 25 ºC under planktonic condition. These results suggested that the metabolic activity of this bacterium was influenced by culture conditions, consequently affecting the subsets of molecules secreted by this strain of *S. marcescens*. Comparison of protein expression profiles between the 25 ºC and 37 ºC static cultures helped us to identify potential proteins with larvicidal activity which include many proteases and chitinases. Protease and chitinase activities were then associated with larvicidal activity against *An. dirus* as supplementation of chitinase or protease inhibitors diminished the larvicidal activity. These proteases and chitinases are likely to damage the larvae at physiological levels rather than at the cellular level because the secreted macromolecules lack cytotoxicity against mosquito cells.

Serralysin, one of the proteases identified from the mass spectrometry analysis of *S. marcescens* secreted macromolecules, has previously been extensively characterized. Serralysin is a metalloprotease first discovered in the culture medium of *Serratia* sp. (Miyata et al., 1970) then later found in other Gram negative bacteria (Baumann, 2013). The protease is secreted as an inactive enzyme and autocatalytically activated in the presence of divalent cations (Baumann, 2013). The enzyme was reported to have entomopathogenic property against larvae of scarab beetle (Pineda-Castellanos et al., 2015). Interestingly, serralysin was found to have immunosuppression property in silk worm and considered as a virulence factor (Ishii et al., 2014). The mechanism of immunosuppression in the study was through the degradation of hemocyte adhesion molecule. Another study demonstrated that a serralysin-type metalloprotease could cleave other proteins involved in insect immunity found in *Manduca sexta* hemolymph (Felföldi et al., 2009). The immune suppression by serralysin could be one of the underlying mechanisms by which *An. dirus* larvae were rendered more susceptible to bacterial infection.

One of the proteases that was detected in our LC-MS analysis is Peptidase M60 enhancin protein, which is a metalloprotease that requires zinc for its function. Enhancin was first described in baculovirus (Hashimoto et al., 1991) then later found in several bacteria and fungi (Nakjang et al., 2012; Slavicek, 2012). Mucin is an important protein component of insect peritrophic matrix including *Anopheles* mosquito (Dinglasan et al., 2009). Enhancin from baculoviruses could degrade intestinal mucin of the insects thus making them more susceptible to viral infection (Toprak et al., 2012; Wang and Granados, 1997) and Bt toxin (Granados et al., 2001). Bacterial enhancin could also digest mucin (Nakjang et al., 2012), but the impact on baculovirus and Bt toxin toxicity was inconclusive (Fang et al., 2009; Galloway et al., 2005; Zhao et al., 2011).

Our results indicate that chitinase and protease inhibitors could only partially reduce the larvicidal activity of the spent medium against *An. dirus* larvae. This suggests that the effect was likely to be a result from a combination of both chitinase and protease enzymes. Although larvicidal activity caused by macromolecules was further reduced when protease inhibitors and chitinase inhibitors were used in combination, the combinatorial effect was marginal and could not completely inhibited the larvicidal activity. Therefore, it is possible that macromolecules other than proteases and chitinases also contributed to the larvicidal activity. This could also be explained by the incomplete inhibition of proteases and chitinases by the inhibitors. As discussed above, several previous studies suggested that larvicidal activity of the identified enzymes were marginal on their own. Contribution of each individual enzyme on larvicidal activity should therefore be further investigated. This can be done by heterologous expression of each enzyme to determine lethal concentrations individually and in combination.

Chitin, a polymer of *N*-acetylglucosamine, is the major component of mosquito extracellular structures including the cuticle of the exoskeleton and the peritrophic matrix (PM) of the midgut in arthropods (Liu et al., 2019). These exoskeletal structures provide strength to the body structure, and serve as muscle attachment sites which allow locomotion and flight (Liu et al., 2019). They are also important barriers for the protection of arthropods from the surrounding environment, mechanical damage, heat, desiccation, and invasion of microorganisms (Liu et al., 2019). With all these important functions, damage to the exoskeleton can be fatal to mosquitoes. Several techniques have been developed to compromise the insects’ exoskeleton including inhibition of chitin synthesis (Belinato et al., 2013; Doucet and Retnakaran, 2012; Zhang et al., 2010), and degradation of the chitin structure (Berini et al., 2019; Singh and Arya, 2019). The later strategy has been exploited using chitinase enzymes or chitinase-producing microorganisms including those from *S. marcescens* (Aggarwal et al., 2015; Binod et al., 2007). However, many studies suggested that chitinases might not be ideal to be used as a stand-alone insecticide for some insects because the larvicidal activity from chitinase can be marginal (Bahar et al., 2012; Zhong et al., 2015). Rather, the enzymes could be used to enhance the insecticidal activity of other control agents including Bt toxin, entomopathogenic virus and chemical insecticides (Kelkenberg et al., 2015; Ramírez-Suero et al., 2011; Zhong et al., 2015). Previous research has demonstrated that chitinases can also help overcome any resistance phenotype developed against a biocontrol strategy. Cai et al. showed that *Bacillus sphaericus* (Bs) toxin-resistant *Culex quinquefasciatus* became susceptible to the Bs treatment when Chitinase AC was overexpressed in *B. sphaericus* (Cai et al., 2007). In addition to the insecticidal property, chitinases can also inhibit fungus by destroying chitinous cell wall structure (Berini et al., 2017; Singh and Arya, 2019; Zarei et al., 2011). *S. marcescens* has been tested to be used against plant fungal pathogen due to its fungicidal properties (Purkayastha et al., 2018) making it an interesting candidate to be used against several classes of agricultural diseases.

Peritrophic matrix (PM) is an acellular structure inside the mosquito gut that compartmentalize gut content from the midgut epithelium (Richards and Richards, 1977; Terra, 2001). It serves as a barrier to protect the insect from the digestive enzymes, abrasion from food particles, and invading parasites (Terra, 2001). PM structure consist of chitin and proteins network, with peritrophins as major protein components (Terra, 2001). Several previous studies have shown the insecticidal activity of microbial chitinases and proteases, and their ability to degrade the exoskeleton and perithrophic matrix (Berini et al., 2019, 2018; Chang et al., 2013; Li et al., 2008; Singh and Arya, 2019). Thus, the larvicidal acticity of our *Serratia* culture-derived fractions in combination to the reduction of larvicidal activity when exposed to chitinase and protease inhibitors hints that this is one if the underlying mechanisms of larval killing. However, more histological-based studies would be needed to concretely confirm this assertion. In addition to direct killing, the compromised peritrophic matrix can be responsible for enhanced susceptibility to bacterial infection observed in our study. For instance, in *Drosophila*, disruption of the peritrophic matrix formation renders the fruit fly more susceptible to bacterial infection (Buchon et al., 2013). The disruption of PM may be the underlying mechanism of the increased larvae’s susceptibility to *Pseudomonas* sp. after *S. marcescens* spent medium treatment.

Since chitin and protein are fundamental building blocks of several organisms, the secreted larvicidal chitinase and proteases can have unintended effects to non-target organisms in the same ecological niches. When chitinases are used for agricultural insect pest management, the enzymes can have additional benefit in fungal disease control as chitin also serve as exoskeleton of fungi (Singh and Arya, 2019). However, *An. dirus* used in this study reside in forestry areas (Obsomer et al., 2007). The application of non-specific chitinase and protease larvicides in such area can affect biodiversity in the forestry environment. On the contrary to the *An. dirus* in Southeast Asia, several reports found adaptation of *Anopheline* mosquitoes in urban environment in Africa (Mattah et al., 2017; Takken and Lindsay, 2019). These urban *Anopheline* mosquitoes might be better targets for the protease- or chitinase-based insecticide due to less potential impact on biodiversity but the efficiency of larvicidal activity has to be confirmed in these species. The shorter duration of the larvicidal effect from enzymatic-based larvicidal molecules suggests environmentally friendly alternative to long-lasting chemical pesticides (Winding et al., 2004).

In summary, this study has identified a strain of *S. marcescens* from a larval rearing habitat in the insectary, that secretes chitinases and proteases that are pathogenic to *An. dirus* larvae. In addition to their intrinsic toxicity, these molecules have potential to increase the larvae’s susceptibility to natural entomopathogenic microorganisms, and other mosquito control strategies currently in use. Although the secreted larvicidal proteases and chitinases have potential for *Anopheles* mosquito control, further studies on concentration optimization, formulation of enzyme mixtures, and enzyme modifications to enhance heat stability are needed for its development into field applications.

## Author contributions

**Natapong Jupatanakul** Conceptualization, Methodology, Validation, Formal analysis, Investigation, Resources, Writing - Original Draft, Writing - Review & Editing, Visualization, Supervision, Project administration, Funding acquisition

**Jutharat Pengon** Methodology, Validation, Formal analysis, Investigation, Writing - Review & Editing

**Shiela Marie Gines Selisana** Methodology, Validation, Formal analysis, Investigation, Writing - Review & Editing

**Waeowalee Choksawangkarn** Methodology, Validation, Formal analysis, Investigation, Writing - Original Draft, Writing - Review & Editing

**Nongluck Jaito** Methodology, Validation, Formal analysis, Investigation, Writing - Original Draft

**Atiporn Saeung** Methodology, Resources, Writing - Review & Editing

**Ratchanu Bunyong** Investigation

**Navaporn Posayapisit** Investigation

**Khrongkhwan Thammatinna** Investigation

**Nuttiya Kalpongnukul** Investigation

**Kittipat Aupalee** Investigation

**Trairak Pisitkun** Methodology, Resources,

**Sumalee Kamchonwongpaisan** Resources, Writing - Review & Editing

## Acknowledgements

We thank Jose Luis Ramirez (USDA) for the suggestions and editorial help with the manuscript. We are grateful to Asst. Prof. Dr. Patchara Sriwichai from Department of Medical Entomology, Faculty of Tropical Medicine, Mahidol University, who kindly provided *An. dirus* adult used to establish the colonies used in this study to A. Saeung.

---

**Supplementary Table 1.**
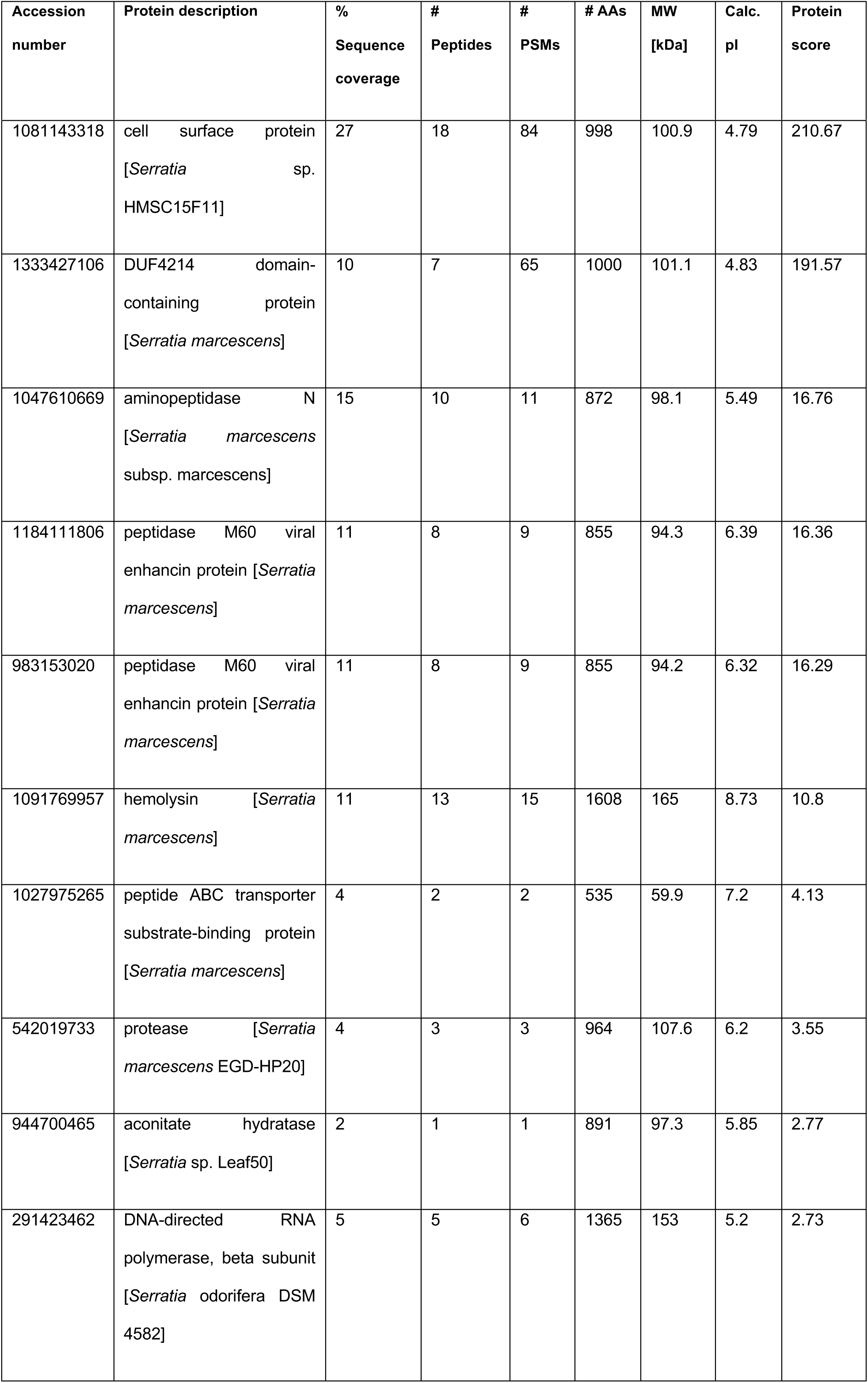
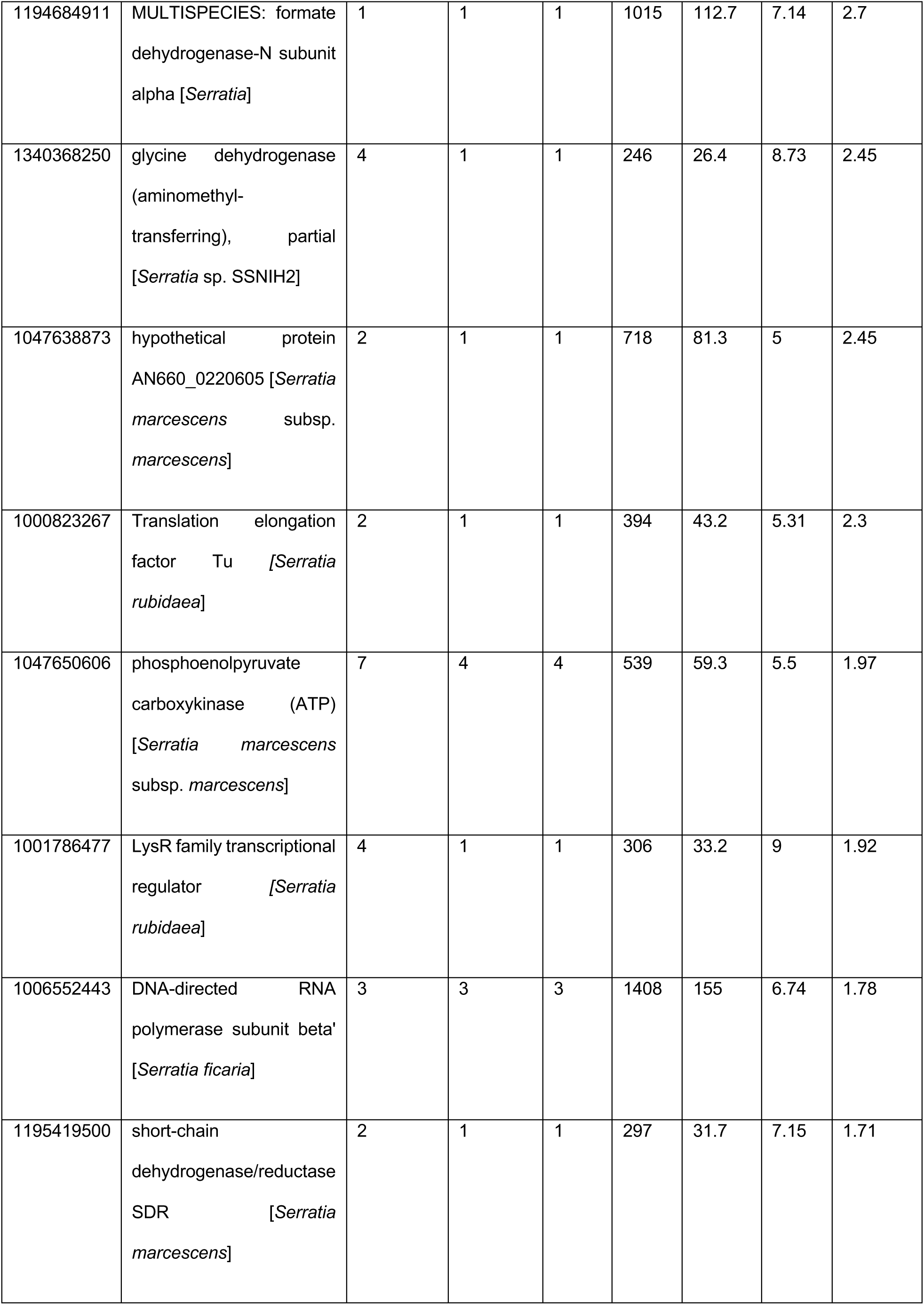
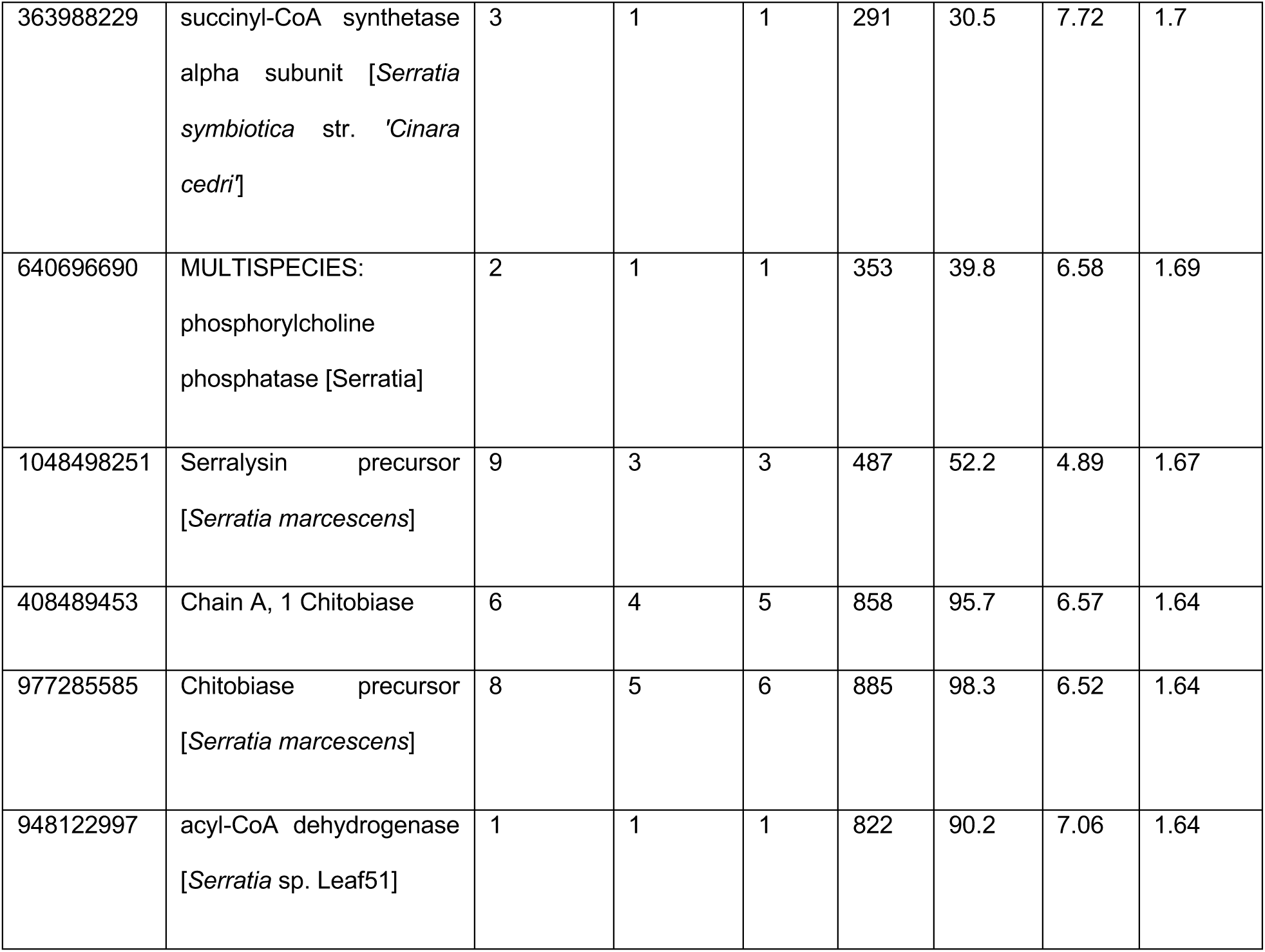
LC-MS analysis of 110 kDa protein band showing the NCBI accession number, protein description, percentage of the protein sequence covered by identified peptides (% sequence coverage), the number of identified peptides (# Peptides), total number of identified peptide spectra matched for each identified protein (#PSM), number of amino acid residues in each identified protein (#AA), molecular weight of proteins (MW), calculated pI of proteins (Calc. pI) and the identification score of proteins calculated by Sequest HT algorithms (Protein score).

**Supplementary Table 2.** LC-MS analysis of 55 kDa protein band showing the NCBI accession number, protein description, percentage of the protein sequence covered by identified peptides (% sequence coverage), the number of identified peptides (# Peptides), total number of identified peptide spectra matched for each identified protein (#PSM), number of amino acid residues in each identified protein (#AA), molecular weight of proteins (MW), calculated pI of proteins (Calc. pI) and the identification score of proteins calculated by Sequest HT algorithms (Protein score).

| Accession<br>number | Protein description | %<br>Sequence<br>coverage | #<br>Peptides | #<br>PSMs | #<br>AAs | MW<br>[kDa] | Calc.<br>pI | Protein<br>score |
| --- | --- | --- | --- | --- | --- | --- | --- | --- |
| 1048498251 | Serralysin precursor [*Serratia*<br>*marcescens*] | 36 | 15 | 76 | 487 | 52.2 | 4.89 | 166.7 |
| 1205048185 | serine 3-dehydrogenase<br>[*Serratia nematodiphila*] | 31 | 13 | 43 | 472 | 50.4 | 4.75 | 96.43 |
| 452850059 | insecticidal serralysin-like<br>protein, partial [*Serratia*<br>*marcescens*] | 52 | 14 | 69 | 320 | 35 | 5.33 | 144.33 |
| 1333426919 | serine 3-dehydrogenase<br>[*Serratia marcescens*] | 25 | 12 | 31 | 472 | 50.4 | 4.74 | 71.38 |
| 1268850421 | serine 3-dehydrogenase<br>[*Serratia* sp. BW106] | 18 | 8 | 29 | 487 | 51.9 | 4.82 | 67.01 |
| 1122396880 | DUF4214 domain-containing<br>protein [*Serratia marcescens*] | 4 | 3 | 12 | 100<br>0 | 101 | 4.87 | 39.48 |
| 1047617935 | arginine--tRNA ligase<br>[*Serratia marcescens* subsp.<br>*marcescens*] | 22 | 12 | 12 | 576 | 63.9 | 5.55 | 18.71 |
| 1039033801 | succinylarginine dihydrolase<br>[*Serratia marcescens*] | 27 | 12 | 12 | 446 | 49.4 | 6.77 | 11.96 |
| 1081143318 | cell surface protein [*Serratia*<br>sp. HMSC15F11] | 10 | 6 | 8 | 998 | 100.9 | 4.79 | 7.17 |
| 1334464241 | histidine ammonia-lyase<br>[*Serratia marcescens*] | 19 | 7 | 8 | 514 | 54.9 | 6 | 12.68 |
| 1127886605 | Proline dipeptidase [*Serratia*<br>*marcescens* SMB2099] | 18 | 7 | 9 | 444 | 50.2 | 6.38 | 11.32 |
| 977423541 | Extracellular serine protease<br>precursor [*Serratia*<br>*marcescens*] | 12 | 8 | 9 | 103<br>4 | 107.8 | 5.39 | 4.24 |
| 1280179082 | capsule assembly Wzi family<br>protein [*Serratia* sp. TKO39] | 17 | 6 | 6 | 478 | 52.7 | 5.41 | 6.61 |
| 1027970958 | Xaa-Pro aminopeptidase<br>[*Serratia marcescens*] | 11 | 4 | 5 | 438 | 49 | 5.38 | 7.85 |
| 1027968317 | glutathione reductase<br>[*Serratia marcescens*] | 16 | 5 | 5 | 450 | 48.5 | 6.38 | 6.32 |
| 1001783318 | phosphoenolpyruvate<br>carboxykinase (ATP)<br>[*Serratia rubidaea*] | 5 | 2 | 2 | 539 | 59 | 5.25 | 1.95 |
| 1216656901 | flagellin [Serratia<br>marcescens] | 11 | 4 | 5 | 423 | 44.4 | 5.24 | 1.99 |
| 493760228 | F0F1 ATP synthase subunit<br>beta [Serratia symbiotica] | 7 | 2 | 2 | 460 | 50.2 | 5.12 | 2.55 |
| 1141208419 | serine 3-dehydrogenase<br>[*Serratia oryzae*] | 6 | 3 | 7 | 522 | 55.7 | 4.54 | 11.77 |
| 1028171175 | hypothetical protein [Serratia<br>sp. YD25] | 4 | 2 | 2 | 568 | 60.8 | 4.65 | 2.17 |
| 676309841 | N-succinylglutamate 5-semialdehyde<br>dehydrogenase [Serratia sp.<br>SCBI] | 2 | 1 | 1 | 639 | 68.5 | 9.8 | 2.13 |
| 383294472 | glucose-6-phosphate<br>isomerase [Serratia sp.<br>M24T3] | 4 | 2 | 2 | 549 | 61.2 | 6.09 | 1.64 |
| 1000823334 | Secreted alkaline<br>metalloproteinase [Serratia<br>rubidaea] | 4 | 2 | 6 | 497 | 53.2 | 4.7 | 9.45 |
| 1057964136 | chitinase [Serratia sp. YD25] | 3 | 2 | 2 | 480 | 51.8 | 5.62 | 2 |
| 1195419500 | short-chain<br>dehydrogenase/reductase<br>SDR [Serratia marcescens] | 2 | 1 | 3 | 297 | 31.7 | 7.15 | 5.5 |
| 1142792582 | serine 3-dehydrogenase<br>[Serratia sp. S119] | 3 | 1 | 1 | 479 | 52 | 4.81 | 1.8 |
| 1006546552 | NADP-dependent isocitrate dehydrogenase [Serratia ficaria] | 3 | 1 | 1 | 417 | 45.6 | 5.68 | 2.03 |

